# Beyond the Cut: Long-read sequencing reveals complex genomic and transcriptomic changes in AAV-CRISPR therapy for Duchenne Muscular Dystrophy

**DOI:** 10.1101/2025.08.01.668007

**Authors:** Mary S. Jia, Made Harumi Padmaswari, Landon A. Burcham, Shilpi Agrawal, Gabrielle N. Bulliard, Abbey L. Stokes, Christopher E. Nelson

## Abstract

Adeno associated virus (AAV)-mediated delivery of CRISPR associated nucleases (AAV-CRISPR) is a promising solution to treat genetic diseases such as Duchenne Muscular Dystrophy (DMD) and is now in early clinical trials. However, genotoxicity and immunogenicity concerns have hindered clinical translation. Due to the complex etiology associated with DMD, the post-transduction consequences of double-stranded breaks induced by AAV-CRISPR in disease models are unclear. This barrier is partially conferred by conventional sequencing methods where common outcomes of AAV-CRISPR editing often escape detection. However, recent reports of novel long-read sequencing approaches permit comprehensive variant detection using a broader sequence context. Here, we comprehensively investigated genomic and transcriptomic post-AAV-CRISPR transduction consequences in myoblast cells and a DMD mouse model following intramuscular and intravenous AAV-CRISPR therapy using both long- and short-read sequencing techniques. Structural variant characterization indicates that unintended on-target large insertions and inversions are common editing outcomes. We demonstrate that combining adaptive sampling with nanopore Cas9-targeted sequencing (AS-nCATS) for long-read quantification of AAV integration is synergistic for detecting difficult-to-amplify editing events. This unbiased data suggests that full-length AAV integration is equally as probable as the on-target deletion. Further, we develop a Nanopore Rapid Amplification of cDNA Ends (nRACE-seq) pipeline for long-read detection of unknown 5’ or 3’ ends of edited transcripts. The nRACE-seq approach effectively detects the presence of AAV-*Dmd* chimeric transcripts, erroneous splicing events, and off-target AAV integration sites. In summary, our findings offer insights into the adaptation of AAV-CRISPR DSB-mediated therapeutics for monogenic diseases and promote the standardization of CRISPR evaluation. We highlight the importance of coupling polymerase-based and polymerase-free methods in long-read sequencing to assess editing outcomes as the field progresses toward clinical applications.

## Introduction

Duchenne muscular dystrophy (DMD) is an X-linked muscle-wasting disease caused by mutations in the *Dmd* gene resulting in the absence of functional dystrophin^1^. While recent advancements in therapeutic approaches, including the first FDA-approved DMD gene therapy, have successfully restored partial dystrophin function, no curative strategies are currently available^2–4^. CRISPR-mediated gene editing holds significant promise as a curative treatment for DMD by targeting and correcting mutations resulting in permanent functional dystrophin restoration^5,6^. The most widely adopted CRISPR therapeutic approach for DMD involves using a double-strand break (DSB) to either delete mutated exon(s) or disrupt splice sites^6,7^. For delivery of Cas9 and target gRNAs, these approaches use adeno-associated virus (AAV) to reach skeletal and cardiac muscle. The combination of AAV delivery and CRISPR-mediated DSBs (here denoted as AAV-CRISPR) for DMD has demonstrated promising therapeutic potential in various disease models^8–10^ and is now in the first clinical trial (NCT06594094). Despite this success, there is a gap in understanding the genomic and transcriptomic consequences of nuclease-based editing for muscle disease.

Post-DSB outcomes, particularly in gene knockout applications, have raised concerns due to their unpredictable effects on various genes and chromosomes^11–14^. Moreover, the combination of AAV and CRISPR-mediated DSBs raises further concerns as reports have documented AAV genome integration at DSB sites into host genomes^15–20^. Currently, PCR-based short-read sequencing at cut sites is the most common method for evaluating CRISPR editing. While valuable owing to its high throughput, this approach may overlook potentially deleterious variations in post-editing outcomes^21^. Recent advancements in sequencing technology now enable a more precise and comprehensive evaluation of the impact of CRISPR gene editing. The increased accuracy of single-molecule long-read sequencing in recent years has driven a shift toward its use, as long-read sequencing can sequence large DNA fragments, facilitating the detection of large structural variants (SVs), complex rearrangements, and full-length vector integration events^18,22,23^.

Sequencing and categorizing diverse CRISPR-mediated editing events present a challenge, however, the complex structure of the *DMD* gene introduces additional challenges for sequencing and analysis. These challenges arise primarily from the AT-rich and repetitive sequences surrounding the mutated regions, which can complicate amplification and introduce bias, increasing sequencing error rates^24–26^. Additionally, the impact of diverse editing outcomes on the *DMD* transcriptome is often overlooked, despite many CRISPR-DMD approaches involving splice site disruptions. Notably, studies have yet to evaluate the impact of SVs or vector integration at the transcriptome level. This highlights the need for more comprehensive sequencing evaluation pipelines to accurately capture the full spectrum of *DMD* editing outcomes at the genomic and transcriptomic levels.

In this study, we evaluated AAV integration at the genomic and transcriptome level using orthogonal sequencing approaches on the AAV-CRISPR-treated DMD mouse model, *mdx*. We assessed the gene-editing outcomes of dual gRNA deletions targeting exon 23 *in vitro* and *in vivo* and used unbiased enrichment strategies to determine the scale and impact of structural variants and unintended AAV integration. Our sequencing strategies revealed a high percentage of AAV integration at the gDNA level and the detection of novel AAV-*Dmd* chimeric transcripts. We were further able to characterize heterogeneous transcription events of AAV expressing Cas9 and gRNAs *in vivo*. These data present significant safety considerations for the general therapeutic strategy of AAV-CRISPR.

## Results

### Tn5-based tagmentation sequencing strategies demonstrate enrichment issues

We generated edits in a DMD mouse model using AAV-encapsulated vectors expressing SaCas9 and guide RNAs (gRNAs). We focused on AAV8 and AAV9 vectors in separate *in vivo* assays due to their inherent specificity for muscle. We used a previously validated pair of gRNAs targeting *Dmd* introns 22 and 23 to excise exon 23, which contains a nonsense mutation in the *mdx* mouse model^8^. The activity of these gRNAs was first confirmed in mouse myoblast C2C12 cells via electroporation and AAV9 transduction using the surveyor assay and deletion PCR. The validated gRNAs were then co-packaged with SaCas9 into both AAV8 and AAV9 vectors and injected intramuscularly into the tibialis anterior (TA) muscle or systemically via tail vein injection in *mdx* mice (**Fig. 1A**). AAV8 constructs carried the gRNAs on a separate vector genome from SaCas9, while AAV9 vectors used a design with Cas9 and a single gRNA in the same vector.

**Figure 1.**
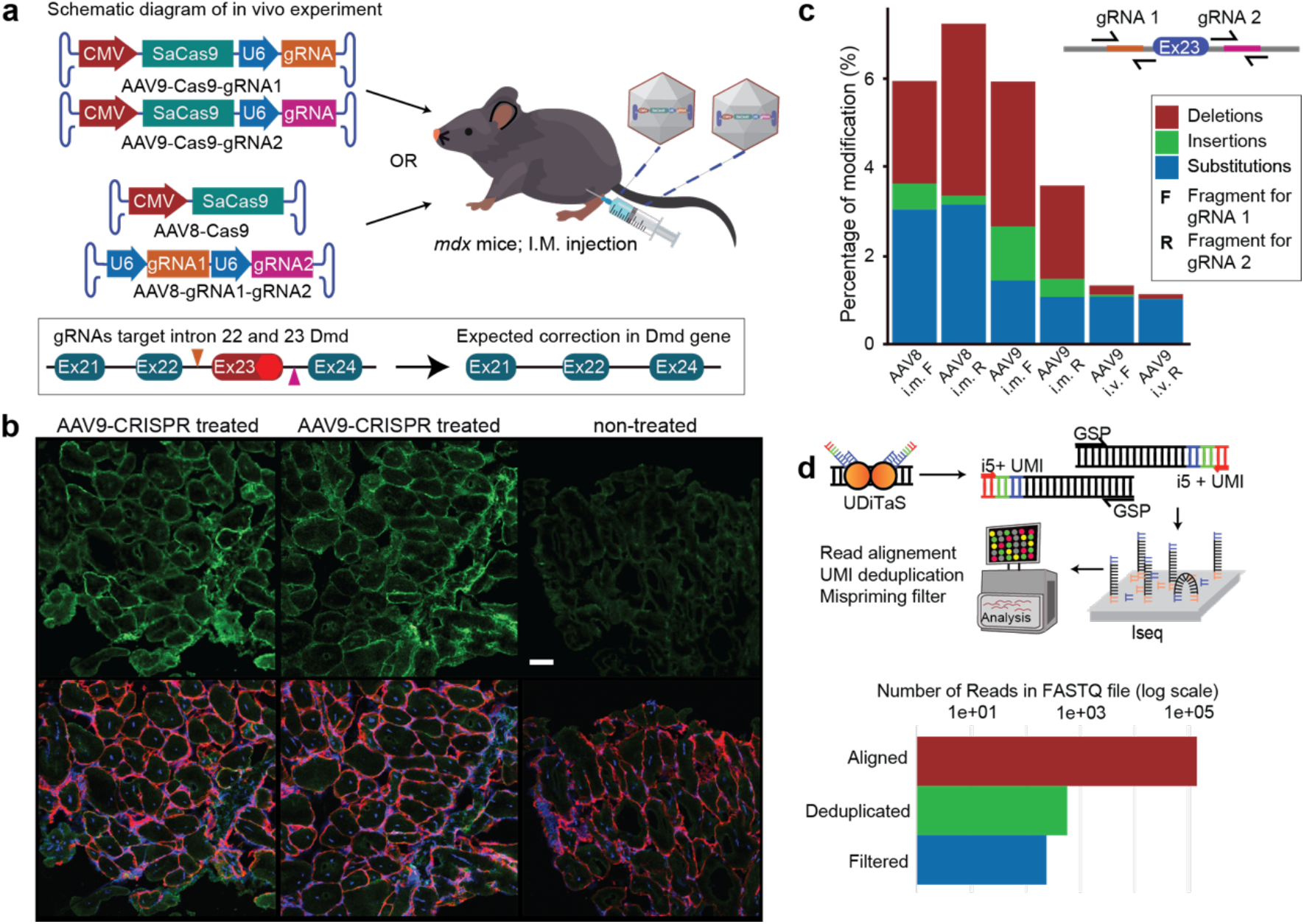
Short-read editing detection strategies. **A)** A schematic of the experimental design for exon 23 deletion using a pair of gRNAs within intron 22 and 23, respectively. The arrows above the target site indicate gRNA targets. Downstream analysis conducted through PCR-enriched short read sequencing was used to verify successful editing and to analyze the range of unintended on-target editing outcomes detectable using short read technologies. **B)** Dystrophin immunofluorescence staining (top) and combined laminin, dystrophin, and DAPI staining (bottom) show dystrophin recovery in the AAV9-CRISPR treated group compared to non-treated. **C)** PCR-enriched editing quantification shows the percentage of modifications detected in the editing window of gRNA target sites using CRISPResso analysis software. Target site F is in intron 22 and R is intron 23. Intravenous reverse guides were excluded as the editing data was indistinguishable from background. **D)** Significant reduction in number of reads in FASTQ file after deduplication and mispriming filter.

Four weeks post-injection, RNA and genomic DNA (gDNA) were harvested from the TA muscle for downstream sequencing analyses. Dystrophin restoration was confirmed by histology for all edited samples (**Fig. 1B**). *In vivo* editing efficiencies were assessed via short amplicon PCR-enriched sequencing of each gRNA target site, revealing similar editing efficiency results for AAV8 and AAV9 delivery in *mdx* mice (**Fig. 1C**). As previously observed, local injections into the TA muscle showed significantly higher editing efficiencies compared to systemic intravenous injections, which had editing rates near wild-type background levels (**Fig. 1C**)^19^. Consequently, we primarily focused on local injection samples to uncover as many editing byproducts as possible.

Using targeted amplicon sequencing to evaluate editing efficiencies introduces the risk of PCR bias, as PCR tends to favor the amplification of high-abundance alleles in a mixed sample. To reduce this bias, we tagged amplicons with Unique Molecular Identifiers (UMIs) during PCR, decreasing the impact of PCR duplicates. To further mitigate biases associated with variable-length PCR amplification and to measure structural changes, we performed UDiTaS Tn5 tagmentation-based enrichment assay (Uni-Directional Targeted Sequencing) and evaluated editing efficiencies using short-read sequencing^27^. However, the coverage of reads was significantly reduced due to the elimination of PCR duplicates, the low efficiency of UDiTaS-based amplification of the i5 adapter, and the false priming of gene-specific primers (**Fig. 1D**). This low coverage makes it challenging to confidently detect large editing outcomes and structural rearrangements at the editing site without significant assay optimization or large-scale sequencing^20^.

### Long-read PCR-enriched sequencing primarily detects large deletions in AAV-CRISPR editing

Most *Dmd* gene editing analyses are performed via short-read sequencing, which is limited to regions immediately up and downstream of on-target and off-target sites, aside from less commonly used tiling strategies^8,19,28^. Despite a high sequencing coverage, this approach overlooks potentially genotoxic editing outcomes resulting from on-target Cas9 cleavage and NHEJ, such as large genomic deletions or rearrangements. Long-read sequencing provides a more comprehensive sequence context, permitting the detection of large structural rearrangements caused by AAV-CRISPR interventions. Using PCR-enriched long-range Nanopore sequencing spanning 10 kb from intron 21 to intron 25, we detected precise on-target deletions both *in vitro* and *in vivo* (**Fig. 2A**). Similarly, sequencing a 3kb amplicon spanning the cDNA up and downstream of exon 23 also resulted in the on-target exon 23 deletion (**Fig. 2B**). Interestingly, a spontaneous exon 27 deletion was also present even in non-treatment, wild-type groups, which we noted for downstream analyses as a possible site for false editing detection (**Fig. 2B**).

**Figure 2.**
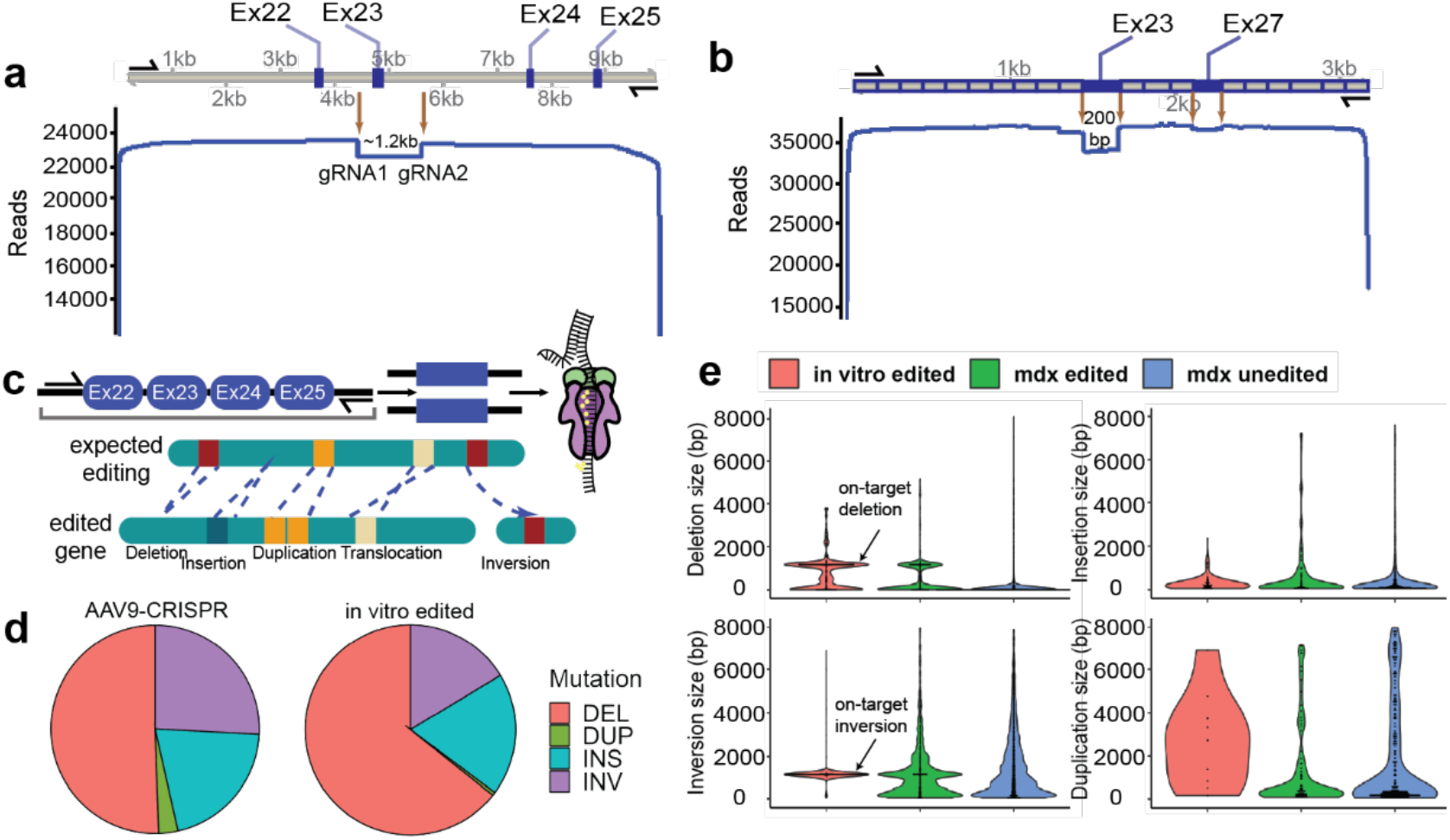
PCR-enriched long-read sequencing detects on-target deletions and inversions. **A)** PCR-enriched long-read sequencing of a 10 kb region shows a deletion in the targeted genomic region. **B)** PCR-enriched long-read sequencing targeting cDNA shows deletion in targeted exon 23 and spontaneous deletion in exon 27. **C)** Illustration of possible unintentional structural variant edits. **D)** A comparison of editing events between *in vivo* AAV9-CRISPR and *in vitro* electroporation. **E)** A size distribution of editing events between *in vivo* AAV9-CRISPR and *in vitro* electroporation.

While short-read experiments categorize any perturbation of a reference sequence as an NHEJ edit, long-read sequencing provides precise resolution of the exact NHEJ editing outcomes, highlighting its advantage in detecting single mutations. As illustrated in **Fig. 2C**, long-read sequencing enables the detection of several distinct types of structural mutations following CRISPR editing. To further characterize these structural variants, sequencing data were aligned to the reference amplicon, and variant detection was performed using the cuteSV and Sniffles pipelines^22,29^. Compared to Sniffles, cuteSV detected a broader range of structural variants due to its reduced coverage requirements (**Fig. S2**). However, both pipelines consistently identified the primary edit *in vivo* as the on-target exon 23 deletion resulting from dual DSBs, with inversions and insertions being the next most common outcomes (**Fig. 2D**).

Notably, in categorizing the number of mutations agnostic of mutation site, all samples including non-treatment illustrate a large number of deletions, insertions, and inversions with few duplication events detected (**Fig. S2E**). These data suggest that a baseline level of sequencing error is present, which would allow us to categorize any significant difference between treated and non-treated samples as coincident with CRISPR editing. A smaller fraction of large SVs include insertion and duplication events that concentrate at the editing site compared to potential sequencing artifacts in non-treatment groups (**Fig. 2E**). This clustering is less evident for insertion events, which appear to have a more stochastic distribution similar to unedited *mdx* mice aside from 3-5 kb AAV integration events (**Fig. 2E**).

### Unbiased Cas9-targeted enrichment and adaptive sampling detect full-length AAV vector integration

While PCR enrichment successfully identified rare structural variant events, PCR bias and complications amplifying repetitive regions such as ITRs compromise reliable editing quantification and full-length AAV detection^18,19^. To reduce PCR-related bias, we initially attempted whole genome sequencing (WGS) with Tn5 library shearing and detected significantly reduced edit-spanning coverage with virtually no editing events (**Fig. S5.B**). While WGS would theoretically capture our editing outcomes of interest, full coverage would require the use of multiple flow cells, which is a significant cost limitation. As an alternative to WGS, previous studies have demonstrated that full-length AAV integration events can be captured using an unbiased CRISPR-enrichment approach^18^. Encouraged by these results, we used nanopore Cas9-targeted sequencing (nCATS) to enrich an approximately 13 kb region spanning the target cut sites. nCATS is an amplification-free library prep method where genomic DNA is dephosphorylated, cleaved at RNA-guided sites flanking the target sequence with Cas9, and adaptor sequences are subsequently ligated to the phosphorylated ends of the cleaved sequence^30^.

We used the CHOPCHOP web tool^31^ to select four gRNA sites flanking our region of interest with high computationally predicted activity and specificity. We performed the original nCATS strategy using these guides, which had high *in vitro* digestion activity (**Fig. S5.A**). Using nCATS, we were unable to successfully detect on-target reads covering the *Dmd* gene. Limited pore occupation due to a depletion of adaptor-ligated sequences resulted in non-detection of our target 13kb region due to a lower sequencing depth than should be achievable considering our high input DNA isolation. To increase the read coverage, we modified the nCATS approach by skipping the dephosphorylation step to increase pore occupation and applied adaptive sampling (nCATS-AS) to reject off-target strands while sequencing in real-time (**Fig. 3A**)^32–34^.

**Figure 3.**
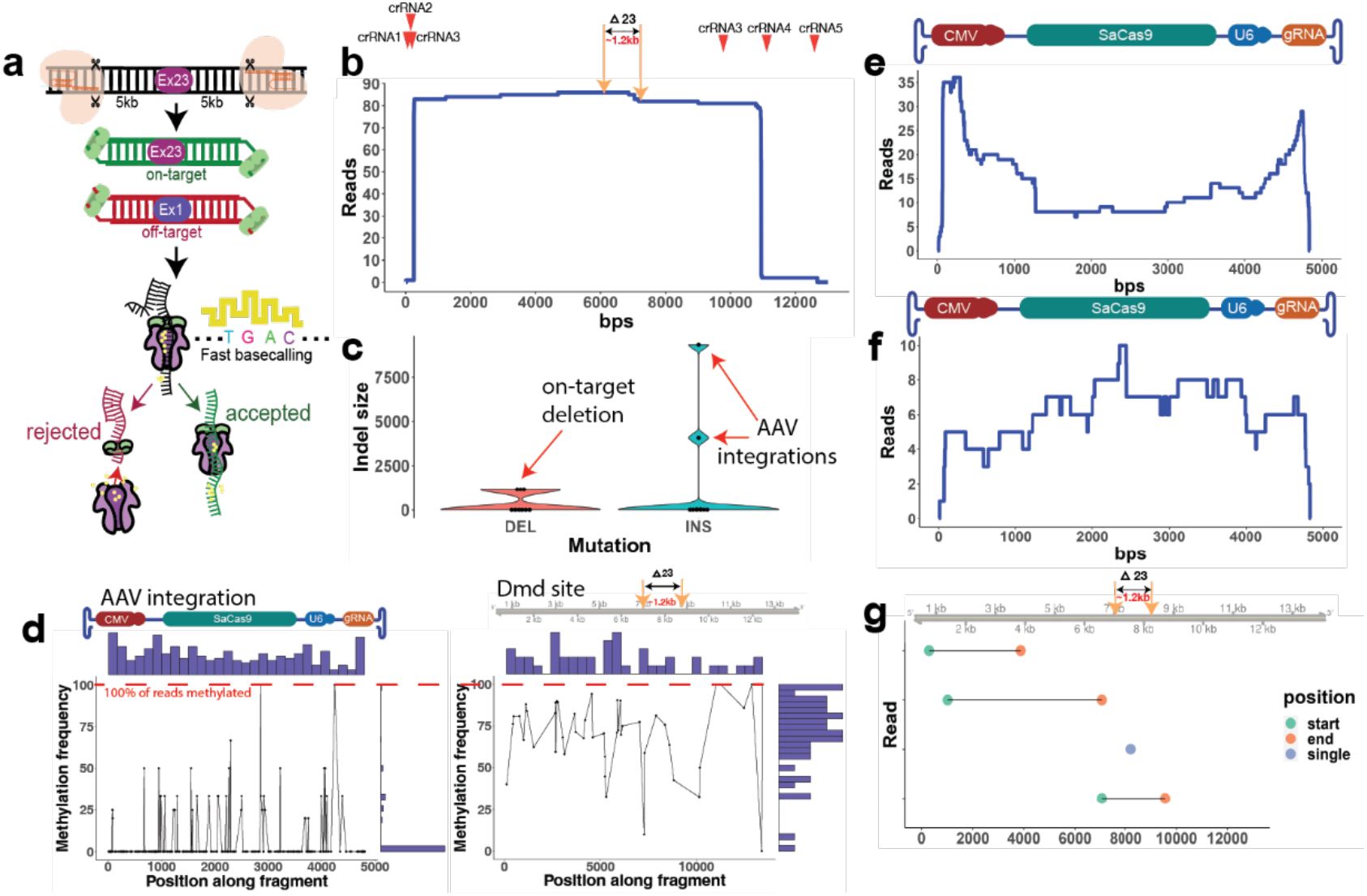
Nanopore Cas9-targeted sequencing and adaptive sampling (nCATS-AS). **A**) Schematic figure of adaptive sampling and nCATS pipeline. **B**) Coverage plot of nCATS in the targeted region using 3 pairs of crRNA, capturing 13 kb. The boundaries of the sequenced reads illustrate that two sets of guides were significantly more efficient cutters despite all pairs demonstrating *in vitro* digestion activity. **C**) Size distribution of insertion and deletion editing events detected by the nCATS method in an in vivo AAV-CRISPR edited sample. **D**) Methylation distribution using bedMethyl annotation of Dorado methylated base-calling algorithms of AAV integrations detected in treated *mdx* and the endogenous *Dmd* site. **E**) Coverage plot of AAV integration from long-read PCR-enriched sequencing. **F**) Coverage plot of AAV integration from the nCATS method. **G**) Mapping of AAV integrations site detected in the *Dmd* sequence demonstrates heterogeneity in integration location.

Adaptive sampling (AS) is an enrichment approach that uses a fast base calling model to reverse voltages, rejecting strands that do not align in real time to a provided reference sequence (**Fig. 3A**). This approach is advantageous as an on-flow cell enrichment strategy selecting for a generally large region of interest to provide increased coverage of an unenriched sample. On-flow cell enrichment enables a high coverage of the target site while avoiding loss of target sites due to off-target depletion in pulldown strategies. By combining enrichment techniques, we achieved 86X coverage of the target region and successfully detected full-length AAV integration between cut sites, reaching up to 4% (**Fig. 3B** and **Fig. 3F**). Notably, the coverage levels as indicated by real-time alignment of nCAT-AS reads were significantly higher (2983x) compared to the final coverage of full-length sequences. When we aligned this aberrant region to the whole mm10 genome sequence, we observed 100% alignment to a fragment on chr 17, indicating an off-target acceptance of repetitive regions in the genome.

Using solely the reads spanning the whole *Dmd* site, we performed structural variant analyses to obtain a quantitative understanding of the fraction of editing outcomes. Interestingly, nCATS-AS structural variant analyses indicated that on-target deletions are just as frequent as on-target AAV integrations (**Fig. S5.C**). The majority of detected AAV integration events were single integration while one event was a head-to-head integration of two tandem AAV genomes at the editing site (**Fig. 3C**). Although integration frequencies of 4% have been corroborated by similar studies^18^, the high frequency of this outcome compared to the on-target deletion has never been observed using PCR-based sequencing methods.

Short-read sequencing of AAV integration has previously been demonstrated to predominantly detect integrations in the ITR region^19^, while our unbiased Tn5 sequencing revealed an almost negligible alignment to fragments of the integrated transgene in both short- and long-read strategies (**Fig. S5.B**). For nanopore long-range PCR enrichment, alignment of the integrated sequence to AAV primarily illustrates integrations at the ITRs with ∼33% of integrations of the AAV8/9 genome being near full-length (**Fig. 3E**). Compared to existing reports of AAV integration profiles with Nanopore amplicon sequencing, we can detect low levels of full-length integration with frequent mutations perturbing the full-length sequence. While nCAT-AS solely identifies full-length integrations, corroborating similar studies using CRISPR-enrichment for AAV integration detection^18^, these integrations similarly contain indels inside the AAV genome. It is equally possible that these mutations in the integrated AAV genome could be a consequence of nanopore sequencing errors, error-prone repair, or errors in AAV formulation.

Using Dorado to analyze the methylation state of the nCAT-AS-derived fragments, we were able to determine that integrated AAV insertions have a significantly lower methylation level compared to the endogenous *Dmd* site (**Fig. 3D**). These data are additionally consistent with reports of lower methylation of AAV integrations, indicating the possibility of sustained endogenous expression of the Cas9 transgene. Using nCAT-AS, AAV integration events were also detected at higher efficiencies compared to the on-target deletion, contradicting the results of PCR enrichment. While PCR-enriched integration sites were primarily mapped to the cut sites with a high fraction of singleton integrations with reads terminating at AAV integrations (**Fig. S3.C**), full-length nCAT-AS integration sites were more stochastically distributed across the target region (**Fig. 3G**).

### RACE-seq reveals *Dmd*-AAV chimeric transcripts

Unintended AAV integrations due to AAV-CRISPR DSBs have been previously characterized, but there are limited studies investigating the impact of integration at the transcriptional level. We previously reported a long-read sequencing strategy to detect unknown 5’ RNA ends using a rapid amplification of cDNA ends (RACE) enrichment technique^35^. To determine the transcriptional consequences of AAV-CRISPR DSBs, we extended this strategy and used 5’ and 3’ RACE to enrich unknown 5’ and 3’ ends of transcripts within *Dmd* transcripts using long-read sequencing of cDNA (**Fig. 4A**). Our initial objective was to determine whether DSB editing resulted in heterogeneous transcriptional outcomes such as isoform switching or early transcript termination. Surprisingly, 5’ RACE using a gene-specific primer in *Dmd* revealed fusion events between *Dmd* and AAV transcripts. When mapped to the AAV genome, the AAV components of these chimeric transcripts predominantly originated from a putative CMV transcriptional start site that differs from the known start site (**Fig. 4B**).

**Figure 4.**
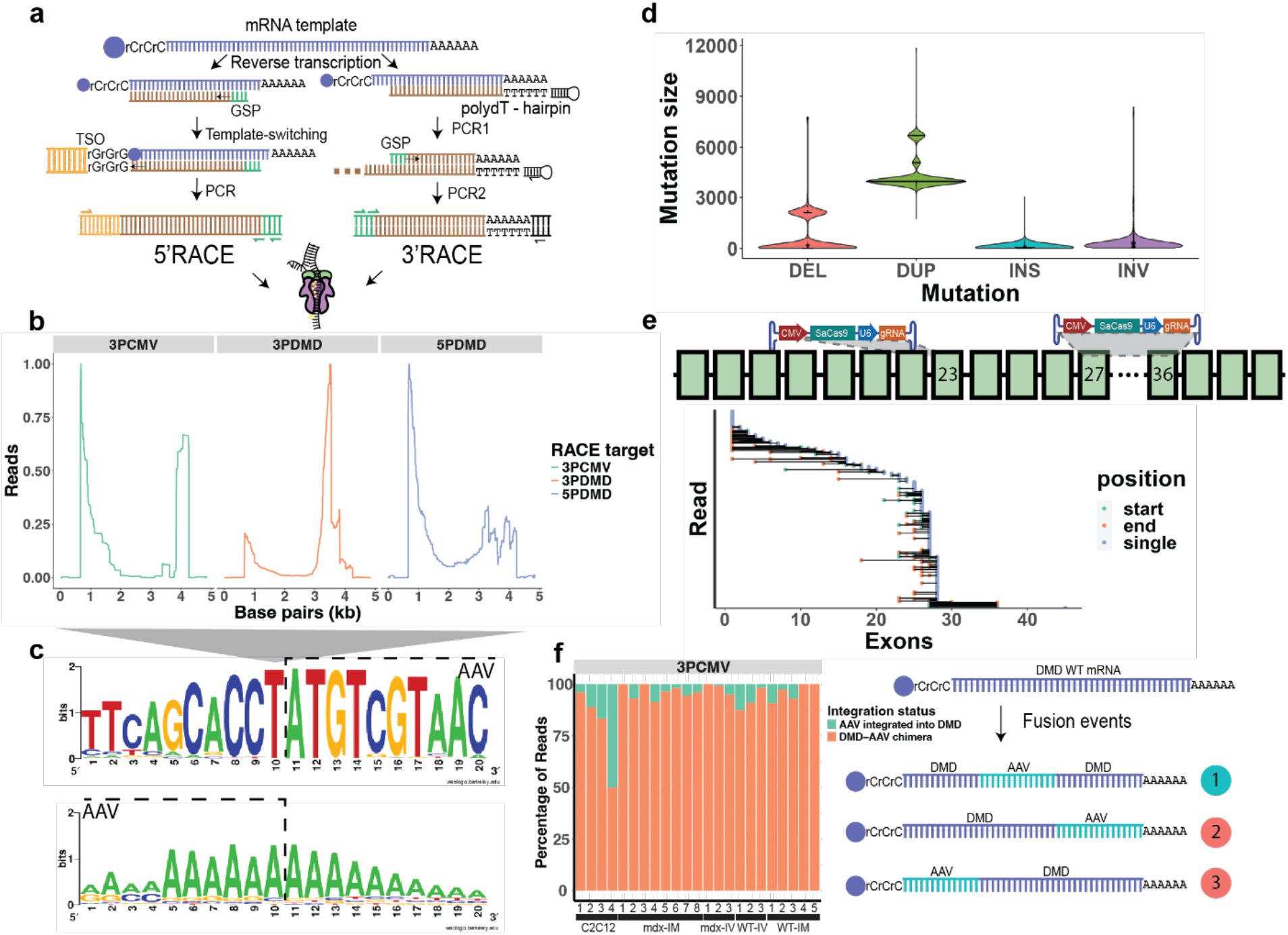
nRACE-seq strategy and variant detection. **A**) Illustration of 5’ and 3’ RACE strategy enrichment for either *Dmd* transcripts or AAV transcripts with details in the methods section. **B**) Coverage of RACE samples in a treated *mdx* TA muscle RNA sample aligned to the AAV genome. **C**) Conserved motifs for integrated *Dmd* transcripts at *Dmd* to AAV transition sites detecting a conserved region present in the CMV promoter sequence and early termination, respectively. **D**) Size distribution of insertion and deletion editing events detected by the RACE method in an in vivo AAV-CRISPR edited sample. **E**) *Dmd* to AAV exon transition sites, illustrating a range of exons removed from the transcript due to unintended integration. **F**) Fraction of reads that terminate/begin with an AAV alignment or reads containing an integrated AAV fragment in 3’RACE enriched samples.

We further analyzed this phenomenon by performing 3’ RACE-seq to determine whether chimeric *Dmd*-AAV transcripts would prematurely terminate in addition to hijacking transcription via the CMV promoter (**Fig. 4A**). We observed 3’ RACE-seq reads terminating early at the bGH poly(A) termination signal (**Fig. S6**), further validating that integrated AAV transcripts can disrupt *Dmd* transcription in addition to expressing SaCas9 and gRNAs. Based on the high mutation rate of integrated AAV genomes as observed both at the gDNA and RNA level (**Fig. 4D, S7** and **S8**), we used RACE-seq enriching for the previously discovered putative CMV transcriptional start site (**Fig. 4B**) and SaCas9 to evaluate the AAV9 transcriptome *in vivo*. Similar to the RACE assays enriching for *Dmd* RNA elements, enrichment of AAV-containing transcripts also detected AAV-*Dmd* fusion events (**Fig. S6**). Notably, a small fraction of non-treated samples illustrate false positive fusion events, which could be used to eliminate false positives from treated samples.

We cross-referenced sequences aligning to both the AAV genome and *Dmd* mRNA to discover the precise location and nature of these fusion events. The dystrophin-AAV fusions discovered in RACE-seq experiments exhibited conserved motifs for fusion mutants with specific integrations that do not terminate or begin with an AAV region (**Fig. 4C**). These conserved motifs suggest cryptic splicing events between *Dmd* and AAV, which result in early bGh signal-mediated termination. Consequently, productive *Dmd* transcripts are likely not produced for these fusion RNA species. We were also able to determine exon drop-off sites at which *Dmd*-AAV chimeric transcripts emerged. Most edited transcripts had integration sites localized to the cut sites from exon 20-25 (**Fig. 4E**). Unexpectedly, a conserved integration site spanning exon 27 was detected at high frequencies in the RACE-seq sequences. This site overlaps with the spontaneous exon 27 skipping site observed in long-read amplicon sequencing of a 3kb cDNA site spanning the edit (**Fig. 2B**). Regions up and downstream of the locations where transcript AAV integration was observed contained splice sites at high frequencies. While this could indicate integration at the gDNA level, it is also possible that these chimeric transcripts could be a consequence of trans-splicing events. While we would be inclined to suspect PCR artifacts were causing these fusion detections, the presence of transcripts that begin as *Dmd*, fuse to AAV, and then terminate with *Dmd* indicates that these results are true fusions (**Fig. 4F**). Interestingly, these integrated fusion transcripts are more frequently detected using 3’RACE enrichments of the AAV genome instead of *Dmd*. This could be due to the background enrichment of unedited *Dmd* mRNA combined with the already rare fusion event.

## Discussion

CRISPR-mediated editing for DMD has been a focus since the advent of CRISPR technology in gene editing. However, there is a lack of well-defined strategies for comprehensive investigations into gene editing outcomes, and no universally accepted “gold standard” method currently exists. This gap highlights the critical need for thorough approaches to analyze gene editing outcomes. In this study, we address this need by employing multiple long-read sequencing techniques to fully examine the outcomes of double-strand breaks (DSBs)—the primary mechanism used in CRISPR-mediated editing for DMD.

Short-read sequencing, specifically PCR-enriched amplicon-based sequencing, stands as the predominant method for assessing the efficacy of CRISPR DMD editing. Its widespread adoption is due to its simplicity and compatibility across various Illumina platforms, including lower throughput systems such as the iSeq100. Despite its ease of use, this technique is susceptible to bias, primarily stemming from the potential overamplification of specific amplicons during PCR^36^. Nevertheless, it remains invaluable for identifying and characterizing the distribution of minor mutations at targeted cleavage sites. To mitigate the biases associated with length-based structural rearrangements detectable via PCR-enriched short-read sequencing, a transposon-based tagmentation technique has emerged^27^. When coupled with UMIs, this approach offers the potential to mitigate amplification biases, enabling quantification at the single-molecule level. However, it entails greater complexity and cost compared to PCR-enriched amplicon sequencing. Moreover, due to the necessity for deeper reads to capture structural rearrangements accurately, this technique requires the usage of higher-end Illumina sequencing platforms^20,27,37^. Commercial library preparation methods like Nextera XT or TruSeq further increase costs, while in-house Tn5 prep reduces expenses but demands greater technical expertise and optimization.

PCR enrichment adaptation to single molecule, long-read sequencing provides an improved method to observe the multifaceted effects of gene editing. The application of long-range PCR enrichment analysis at both gDNA and mRNA levels for DMD has not been previously reported. Using these two methods, we successfully demonstrated a precise deletion of the entire exon 23 at both the gDNA and mRNA levels. This deletion can only be accurately detected at each cut site using short-read sequencing. Additionally, we identified various structural variants and AAV insertions of up to 5 kb. While long-read sequencing expands the sequencing window, PCR error can significantly impact the detection of structural variants (SVs) by introducing false positives or negatives. In our efforts to mitigate these errors, we reduced the cycle number to 15 and used a high-fidelity DNA polymerase. Despite these measures, studies have shown that PCR-mediated base substitution errors still arise from thermocycling-induced DNA damage^38^. Furthermore, other studies reported false-positive results or PCR-chimera using this approach^39,40^. Thus, we determined that additional unbiased sequencing tools are necessary to achieve absolute SV quantification.

To circumvent the use of long-range PCR, Gilpatrick et al. introduced an nCATS enrichment method^30^. This approach has been adapted by others and has demonstrated that Cas9-targeted sequencing generally provides greater coverage of AAV cargo integration, whereas PCR-enriched sequencing predominantly reveals higher coverage of only the ITR regions^18^. The consistent coverage observed between the ITRs across AAV cargos suggests there are no sequence hotspots or specific microhomologies shared between the on-target sites and the AAV cargo. Although Cas9-targeted sequencing achieves higher throughput in terms of AAV integration coverage, the overall enrichment remains poor at most sites^18,30,41,42^. To address this, eliminating the dephosphorylation step and incorporating additional on-board enrichment using adaptive sampling significantly increased coverage compared to the original nCATS protocol. However, the presence of repetitive regions in *Dmd* led to the mis-enrichment of off-target sites, which were subsequently removed during downstream bioinformatics analyses. This highlights the importance of carefully selecting gRNAs outside of repetitive regions in the genome. While we observed significantly improved Cas9-targeting coverage using AS, other studies have shown only moderate improvements in coverage with inconsistent results in achieving higher on-target depth compared to nCATS only^41,43^. This could be because the AS approach rejected most DNA strands, resulting in lower overall throughput compared to using only nCATS. While our findings and those of others reported an increase in the percentage of on-target reads, it is also noted that AS requires high-performance computing resources^32,41,43^.

A comprehensive analysis of structural variants at the transcriptome level following CRISPR editing has been limited, primarily due to the limited technology of long-read RNA sequencing. While the short-read RNA sequencing method is a standard for evaluating transcriptome changes^44^, it falls short in detecting the full spectrum of structural variants. Although we have performed long-range PCR for cDNA amplification, the use of specific forward and reverse primers restricts sequencing to known sites. To achieve a more comprehensive analysis, performing RACE for both the 3’ and 5’ ends would better capture the unknown regions^45,46^. By combining RACE with long-read sequencing, it becomes possible to thoroughly characterize structural variants at the transcriptome level. A recent study by Adamopoulos *et al*. also demonstrated the utility of this approach by combining RACE and nanopore sequencing, which not only successfully mapped the annotated 5’ UTR but also identified novel transcripts in members of the human *KLK* gene family^47^. The nRACE-seq technique we developed successfully detected edited RNA transcripts with the precise deletion expected from exon 23 skipping. Additionally, nRACE-seq revealed the presence of *Dmd*-AAV chimeric transcripts resulting from AAV integration byproducts. These chimeras appeared to cause early transcript termination and initiate transcription via the integrated constitutive CMV promoter. A conserved deletion in exon 27, previously observed during *in vitro* editing, was also detected in the nRACE-seq datasets. This exon demonstrated a high frequency of AAV integration events, indicating that AAV integrations can occur in regions where internal RNA regulation influences exon inclusion or exclusion. The discovery of this novel integration site highlights the importance of using long-read sequencing to uncover broader genome editing byproducts.

## Conclusion

In contrast to reports of low AAV integration rates, long-read sequencing reveals that on-target and off-target AAV integrations are relatively common and can impact the transcription products of the integration sites. We have demonstrated that a high fraction of editing events are AAV integrations and that these integrations can interfere with the intended edited transcript, potentially leading to phenotypic consequences such as neoantigen formation, tumorigenicity, or genotoxic effects. However, our study is not designed to determine the functional impact of these integrations and further studies will be required to assess these risks. With the advent of new gene editing strategies aimed at improving editing precision and reducing reliance on DSBs, robust validation methods are still essential to assess their accuracy and safety^48^. We have additionally introduced a novel sequencing pipeline for unbiased long-read sequencing and enrichment of unknown 3’/5’ end transcripts for characterizing the impact of AAV-CRISPR on RNA. These results provide significant considerations for future applications of AAV-CRISPR technologies on gene therapies.

## Materials and Methods

### AAV Preparation

Guides targeting the mouse *Dmd* gene were selected and cloned into an AAV backbone expressing SaCas9 under the CMV promoter and a sgRNA under the U6 promoter (**Table S1**). Scrambled vectors were also produced in addition to two separate intron 22 and intron 23 targeting constructs as transfer plasmids. ITRs were verified in all transfer plasmids by SmaI digest before production. Each transfer plasmid containing the AAV genome was co-transfected with the rAAV9 or rAAV8 rep/cap plasmid and an adenovirus helper plasmid by the University of North Carolina – Chapel Hill Viral Vector Core and purified using imidazole columns. Titers were provided at 1.5 × 10^13 vg/mL (pCMV-SaCas9-u6-gRNADmd1), 1.8 × 10^13 vg/mL (pCMV-SaCas9-u6-gRNADmd2), 1.7 × 10^13 vg/mL (pCMV-SaCas9-u6-gRNAScr), 2.3 × 10^13 vg/mL (pCMV-SaCas9), 1.2 × 10^13 vg/mL (pU6-gRNAs) dialyzed w/350 mM NaCl & 5% D-Sorbitol in PBS.

### Electroporation and AAV transduction of myoblasts

Plasmids containing SaCas9 and two gRNAs targeting exon 23 were electroporated into C2C12 myoblasts at a 1:1:1 ratio in high glucose DMEM with 20% FBS and 1% pen/strep. Following three days of culturing, the media was replaced with differentiation media comprising 2% equine serum, 1 µM insulin, and 1% pen/strep. The myoblasts underwent differentiation into myotubes for 7 days. Genomic DNA (gDNA) and RNA from the resulting myotubes were extracted using the DNeasy Blood and Tissue kit (Qiagen #69504) and the RNeasy RNA extraction kit (Qiagen #74104), respectively. AAV9 was transduced into C2C12 myoblasts at an MOI of 1: 50, supplemented with 8 ng/µL polybrene. The resulting cells were cultured for three days, followed by a media replacement with differentiation media. Subsequently, gDNA and RNA were extracted using the described methods.

### *In vivo* administration of AAV-CRISPR in *mdx* and wild-type mice

All *in vivo* protocols outlined in this study have received approval from the Institutional Animal Care and Use Committee (IACUC). Wild-type (C57BL/6J) and *mdx* (C57BL/10ScSn-Dmdmdx/J) mice, sourced from Jackson Labs at 8 weeks old, underwent a one-week acclimatization period. Both WT and *mdx* mice received injections either in the TA muscle or via the tail vein with 40 or 200 µL, respectively, of approximately 1E+13 vg/kg of AAV8 or AAV9 for intramuscular and 1E+14 vg/kg for intravenous injection.

For both intramuscular (IM) and intravenous (IV) injections, three mice were assigned to each treatment group, encompassing AAV9 or AAV8-CRISPR targeting intron 22 and 23, AAV9-CRISPR with scrambled guides, and PBS or non-treated as a control. Following a 3-4 week post-injection period, the mice were euthanized, and TA muscles and hearts were collected for subsequent analysis.

### Muscle histology

TA muscle and cardiac muscle sections, embedded in optimal cutting temperature (OCT) compound, were cryosectioned into 10 µm slices on glass slides. Following a triple wash with PBS for 10-15 minutes, the tissue underwent blocking for one hour in a solution consisting of PBS, 5% fetal bovine serum (FBS), 5% goat serum, and 0.5% Triton-X. Immunofluorescence staining was carried out by incubating the slides overnight in a mixture of blocking serum, MANDYS8 (diluted 1:200, Sigma #D8168), and alpha Laminin (diluted 1:1000, Sigma #L9393). After a 24-hour incubation period, tissues were washed thrice with PBS and incubated for an additional hour in blocking serum containing secondary antibodies and 4’,6-diamidino-2-phenylindole (DAPI) (Invitrogen #D3571). Following a three-time rinse with PBS, the slides were mounted using Fluoromount G at room temperature (RT) in darkness for 15 minutes. Imaging was performed using a confocal microscope.

### gDNA and RNA preparation

The TA muscle and heart tissues were processed using the Monarch High Molecular Weight (HMW) DNA Extraction kit from NEB (T3060), following the optimized manufacturer’s protocol for muscle tissue. In summary, skeletal muscle tissues were lysed with a pestle in microtubes and subjected to thermal agitation for 15 minutes, while heart muscle tissues underwent the same process for 45 minutes at 56°C and 2000 rpm. HMW genomic DNAs were then captured by beads and eluted.

For RNA extractions, the TRIzol/chloroform extraction method was employed with the Monarch Total RNA Miniprep Kit (NEB #T2010). Briefly, muscle tissues were homogenized in microtubes using a pestle in 900 µL of TRIzol. After incubating at RT for 5 minutes, 100 µL of gDNA eliminator from the RNAeasy Plus Universal Mini Kit (Qiagen #73404) was added and shaken vigorously for 15 seconds. In the same tube, 180 µL of chloroform was added, shaken vigorously for 15 seconds, and set aside to incubate at RT for 2 minutes. The resulting solution was centrifuged for 15 minutes at 4°C and 12,000g. The clear supernatant was separated without disturbing the white interphase or pink organic layer. An equal volume of 70% ethanol was added to the supernatant and mixed thoroughly by pipetting. The mixture was transferred to an RNA purification column from the Monarch RNA miniprep kit and followed according to the manufacturer’s instructions after the lysis and gDNA elimination step.

### Amplicon sequencing and short-read sequencing

Genomic DNA from TA muscles were PCR-amplified targeting intron 22 and 23 (Table S1). A second PCR round added the Illumina adapter sequence and sample barcodes. PCR products were purified using AMPure beads 1.8X and measured by Qubit. The purified PCR products were pooled equimolarly and sequenced with 150 bp paired-end Illumina iSeq. Demultiplexing using assigned barcodes and trimming of adapter/primer sequences were performed. Crispresso pipeline was used for indel analysis^49^.

### Tn5 Transposon-mediated target enrichment and library preparation

Unloaded Tn5 transposase proteins were obtained from Diagenode #C01070010-10. Tn5 transposase was loaded with oligos containing mosaic ends with 13 base-pair unique molecular identifiers (UMI) and the i5 sequencing adapter following the manufacturer’s protocol. Tagmentation of 50 ng gDNA with 1:8 dilutions of loaded Tn5 was completed as previously described ^27^. Nested PCRs were performed using the i5 adapter and a gene-specific primer up/downstream of the Cas9 cut sites using Q5 Hot Start High-Fidelity DNA Polymerase (#M0491) to enrich the target site and add sample barcodes/the i7 adapter. Sequencing was run on the Illumina iSeq using the 150 bp pair-end reagent cartridge. Alignment of reads was done using bwa-mem2 to the reference genome (DMD amplicon)^50^.

The 13 bp unique molecular identifiers (UMIs) were discovered and collected using the UMI-tools software^51^ and aligned to the target region using bowtie2 short read aligners. The resulting data was converted to a bam and vcf file using samtools and bcftools, respectively. The VCF data was analyzed to determine whether indels were present at gRNA targeting sequences.

### Long-read nanopore amplicon assays

Genomic DNA (gDNA) was amplified using Long Amp PCR with primers targeting the 5 kb region at the 5’ and 3’ cut sites. After PCR, products were purified using 1X AMPure beads and quantified with Qubit. Amplicon libraries were barcoded using Nanopore SQK-LSK109 with EXP-NBD104, following the manufacturer’s protocol. Briefly, amplicon DNA was repaired to achieve 5′ phosphorylated and 3′ dA-tailed ends. DNA was then ligated with sample barcodes and sequencing adapters before sequencing on the MinION platform.

### CRISPR-enrichment and adaptive sampling nanopore sequencing

We ordered crRNAs (**Table S1**) and tracrRNA from IDT and resuspended the RNA oligos to 100 µM in the duplex buffer. Equimolar crRNAs were combined by adding 0.75 µL of each crRNA to a PCR tube. The gRNA duplex was assembled by combining 8 µL of nuclease-free water with 1 µL of 100 µM tracrRNA and 1 µL of the crRNA mix in a PCR tube. The duplex was annealed by incubating at 95°C for 5 minutes and allowing the mixture to cool to room temperature. Ribonucleoproteins were assembled with 23 µL of nuclease-free water, 2.8 µL of 10x CutSmart Buffer (NEB #B7204), 3 µL of the gRNA duplex, and 1.2 µL of the 1:5 Cas9 dilution.

For sequencing runs following the precise nCATS strategy, high molecular weight gDNA was dephosphorylated by adding 3 µL of QuickCIP (NEB #M0525S) and 3 µL of 10X CutSmart Buffer to 3 ug of gDNA in a total volume of 30 µL, and incubating at 37°C for 10 to 30 minutes, deactivating the enzyme by incubating at 80°C for 2 minutes. The resulting dephosphorylated gDNA was cleaved by adding 10 µL of the CRISPR RNPs and monoadenylated with 1 µL of 10mM dATPs and 1 µL of Taq polymerase to the whole 30 µL. We incubated the RNP-gDNA mixture in a thermal cycler at 37°C for 15-30 minutes, 72°C for 5 minutes, and held the reaction at 12°C.

A ligation mix was prepared using 20 µL of LNB from the Nanopore SQK-LSK114 kit, 4.5 µL water, 10 µL of Quick T4 Ligase (NEB #M2200), and 3.5 µL AMX. The ligation mix was added to the Cas9-RNP mix in an Eppendorf tube and was rotated on a hula mixer at RT for 10 minutes until the ligation reaction was complete. An equivolume of TE was added to the ligation reaction and 0.3X Ampure beads were used to purify the adapter-ligated gDNA. The beads were rotated on a hula mixer for 5 minutes to allow the beads to bind and were placed on a magnetic rack for 3 minutes. The supernatant was removed without disturbing the beads, and 200 µL of LFB was used to resuspend the beads with gentle flicking before returning the beads to the magnetic rack. This step was repeated for a second LFB wash. After the second wash, the LFB was fully removed, and beads were eluted in 15 µL of EB at 37°C with gentle agitation for 30 minutes.

For library preps using adaptive sampling, the dephosphorylation step is omitted, but the remaining steps of the preparation are carried out as described. Adaptive sampling was executed using the MinKNOW software by supplying the ∼13 kb region in a fasta format for real-time alignment with fast guppy base calling. Adaptive sampling experiments were conducted on a MinION for 20 hours to achieve high coverage of target sequences.

### 5’ and 3’ RACE nanopore sequencing

All Rapid Amplification of cDNA Ends (RACE) library preps were performed using the Template Switching RT Enzyme Mix (NEB Cat. #M0466S). For 5’ RACE experiments enriching for either the *Dmd* gene or the AAV genome transgenes, a gene-specific primer (GSP1) and a template switching oligo (TSO) were synthesized at IDT with an adapter sequence for amplification and three guanine ribonucleotides (**Table S1**). 1 µL of a 10 uM GSP1 was added to up to 4 µL of an RNA template (10 ng – 1 µg) with 1 µL of 10 mM dNTPs and diluted to a maximum volume of 6 µL. The tube was gently flicked until mixed, spun down, and incubated for 5 mins at 70°C in a thermocycler, and held at 4°C. First-strand synthesis was performed by adding 2.5 µL of the Template Switching RT Buffer (4X) with 0.5 µL of the 75 µM TSO and 1 µL of the RT Enzyme mix (10X) to the previous reaction to a total of 10 µL. After mixing by gentle flicking, the 10 µL reaction was incubated at 42°C for 90 mins, 85°C for 5 mins, and held at 4°C. After a cDNA template was formed, nested PCR was performed using the Q5 High-Fidelity Hot Start enzyme. Up to 2.5 mL of the template-switching cDNA product was combined with 5 uL of the Q5 Reaction Buffer, 0.25 mL of Q5 High-Fidelity Hot Start Polymerase, 1.25 mL of a TSO-specific primer, a second GSP2 downstream of GSP1, and water to a total volume of 25 mL. Touch-down PCR was performed on the resulting reaction using incubation steps as recommended by the manufacturer for 15 cycles. The resulting reaction was purified using 1X AMPure beads, and touch-down PCR was repeated with a GSP3 downstream of GSP2 for an additional 15 cycles. The resulting amplicons were validated for RACE enrichment using qPCR and sequencing samples that demonstrated Cq values lower than that of control samples lacking the gene-specific primer. The average size of the amplicons generated in RACE experiments was determined by gel electrophoresis.

For 3’ RACE sample preps, the same steps were performed using solely a poly(dT) oligo containing an adapter with a hairpin to prevent stochastic binding to internal poly(A) sequences in the first-strand synthesis step. The RT reaction was still performed using the Template Switching RT Enzyme mix using a GSP1 and the annealed poly(dT) oligo to enrich for polyadenylated transcripts containing either an AAV transgene or the Dmd transcript. Barring this step, the same protocol as in 5’ RACE was performed as described using the poly(dT)-oligo-specific adapter sequence rather than a TSO-specific primer in the nested PCR steps. The same validation strategies for 5’RACE-seq were used to validate successful 3’ RACE enrichments. RACE-seq samples were pooled for Nanopore sequencing using the SQK-NBD114.24 kit according to the manufacturer’s instructions using the LFB. Samples were run for 16 hours on a MinION.

### Nanopore sequencing analysis pipelines

The analysis pipeline used for variant calling, methylation detection, and visualization are available at https://github.com/maryjia/NanoporeSVanalysis, including a sample dataset to replicate all analyses presented here. Long-read sequencing data were analyzed using the minimap2 long-read aligners, along with the structural variant detection software cuteSV and Sniffles2^22,29^. Fastq files, demultiplexed using commercial nanopore barcodes, were trimmed and filtered for quality using NanoFilt and then aligned to target amplicons or whole target transcripts/genome sequences^52^. The resulting data were converted to a BAM file using samtools and imported into R to determine the number of reads aligning to the target sequence.

RACE datasets were aligned to the whole *Dmd* or AAV genome. To extract reads up and downstream of *Dmd* and AAV alignments, 20 base pairs up and downstream of aligned regions were extracted and aligned back to the reference using BLAST.

## Acknowledgments

We would like to thank Professor David Ussery for helpful conversations on nanopore sequencing and Professor Dongsheng Duan for sharing gene editing mouse samples. This work was supported by an NIH/NIBIB R00EB023979, NIH/NIGMS R35GM155433, NIH/NIGMS S10OD032338, ASGCT Career Development Award, University of Arkansas Chancellor’s Innovation Grant, and The Arkansas Bioscience Institute. MHP was supported by the Fulbright Indonesia Research Science and Technology (First) Master’s Degree Program and PhRMA Foundation 2023 Predoctoral Fellowship in Drug Discovery. G.B. was supported by the University of Arkansas Honors College Undergraduate Research Grant. M.S.J. was supported by a State Undergraduate Research Fellowship (SURF), the Bodenhamer Fellowship, and the Goldwater Scholarship. CEN was supported by the 21st Century Chair in Biomedical Engineering. AS was supported by the Distinguished Doctoral Fellowship. SA was supported by a Women’s Giving Circle Grant.

