## Supplemental Information for "Beyond the Cut: Long-read sequencing reveals complex genomic and transcriptomic changes in AAV-CRISPR therapy for Duchenne Muscular Dystrophy"

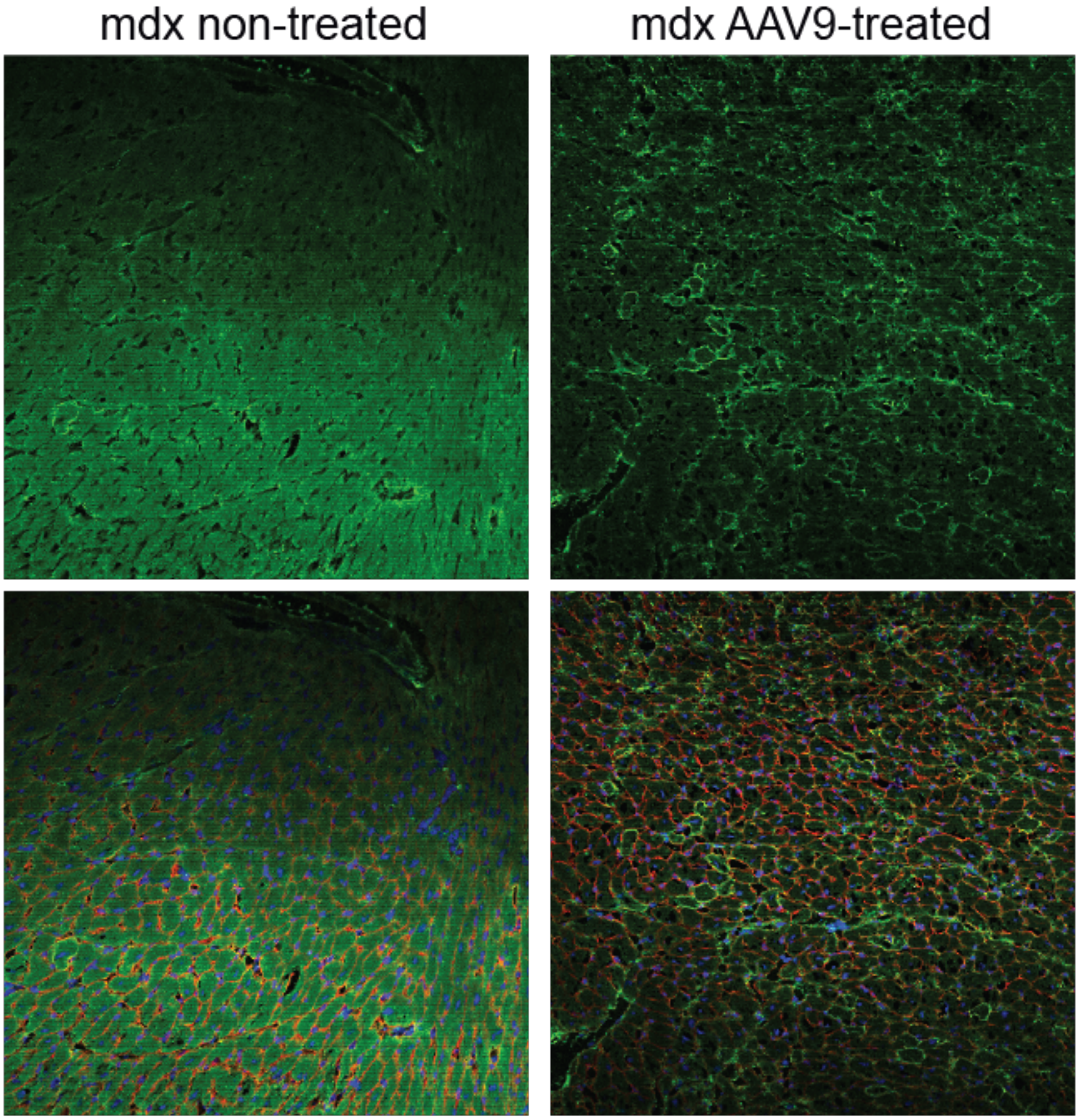


**Figure S1.** Heart histology of non-treated and systemic AAV9 injected *mdx* mice. Demonstrates low editing efficiency in heart tissue compared to skeletal muscle tissue.

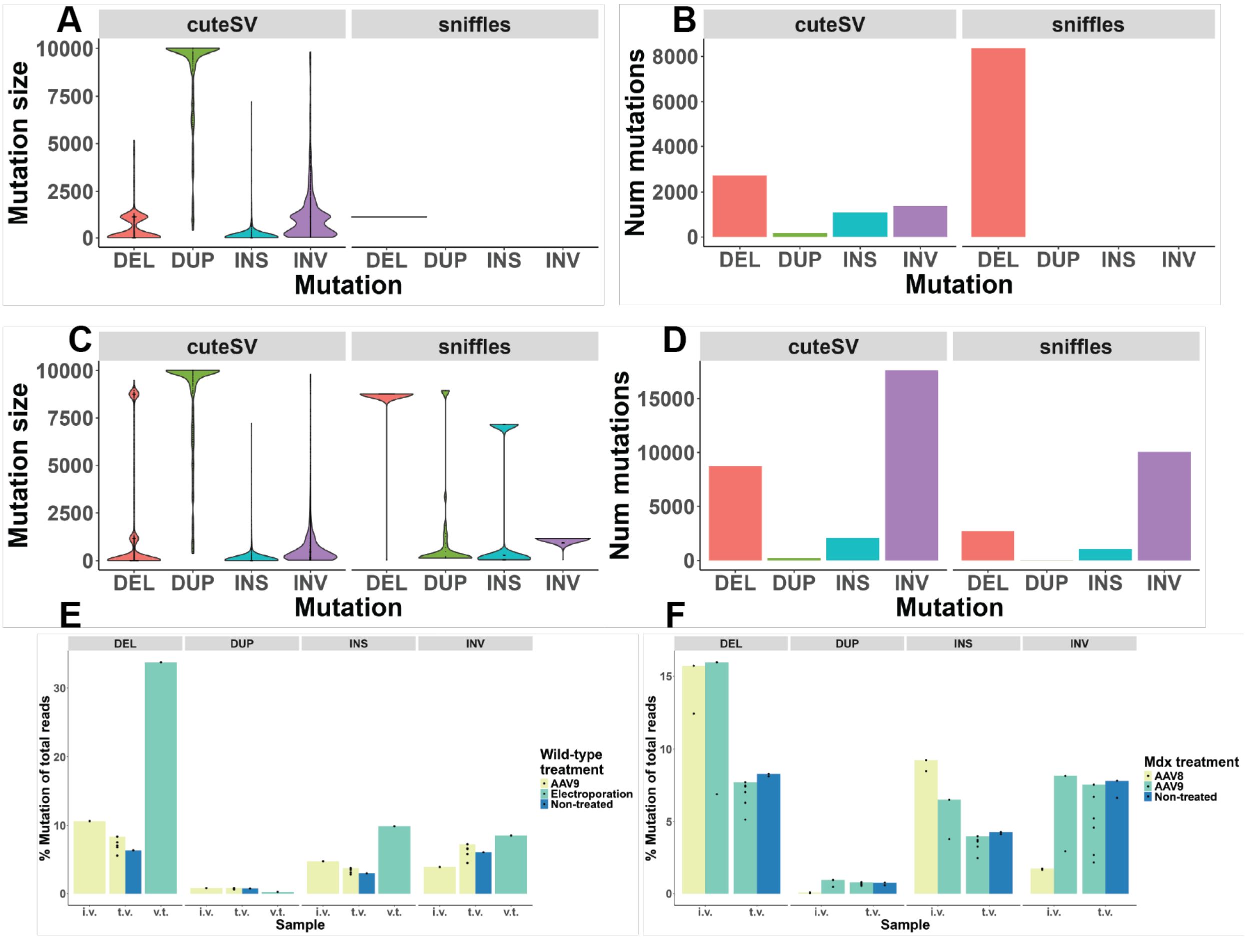


**Figure S2.** Comparison of cuteSV and sniffles structural variant annotation software. The cuteSV software detects a larger number of mutation events and accurately detects the target site deletion compared to sniffles. However, the sniffles pipeline also precisely detects the on-target edits. **A&B)** The more restrictive minimap2 alignment parameters selecting for lengths > 1000 bps and trimming adapters results in sniffles detecting only the precise on-target deletion while cuteSV reports a more diverse range of edits usually centering around the cut site. **C&D)** The flexible alignment strategy results in a similar profile of SV detection across both software, but with sniffles failing to detect the on-target edit. **E&F)** Overall % mutation for all 10 kb sequenced samples over the total number of reads. Sample annotated with t.v. were administered systemically, samples annotated with i.m. were administered intramuscular injections, and the v.t. annotations indicate *in vitro* assays. The sample types were separated by model (e.g. wild-type and *mdx*, respectively).

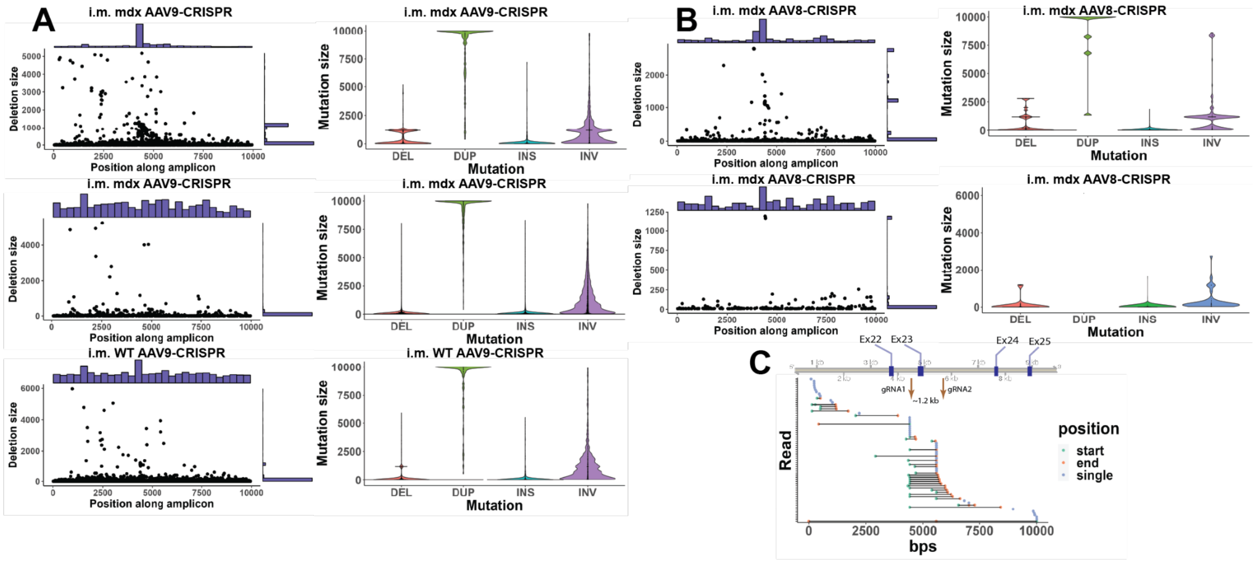

**Figure S3**. Mutation profiles following intravenous AAV9-CRISPR delivery detected via long-range amplicon sequencing. Deletion size and positional distributions of AAV9-CRISPR (**A**) and AAV8-CRISPR (**B**) treatment following intramuscular injection of each respective AAV in either *mdx* or wild-type (WT) mice. The left column displays violin plots showing the distribution and size range of detected mutations, categorized by type: deletions (DEL), duplications (DUP), insertions (INS), and inversions (INV), illustrating a consistently high number of mutation events detected. Each panel corresponds to a biological replicate. These profiles highlight differences in mutation types and their distribution across the target regions in both *mdx* and WT mouse models. **C**) Plotting the position at which AAV integration begins and terminates for amplicon sequencing of one biological replicate of the *mdx* mice treated with AAV9-CRISPR demonstrates a clustering of integration events at the location of gRNA targets.

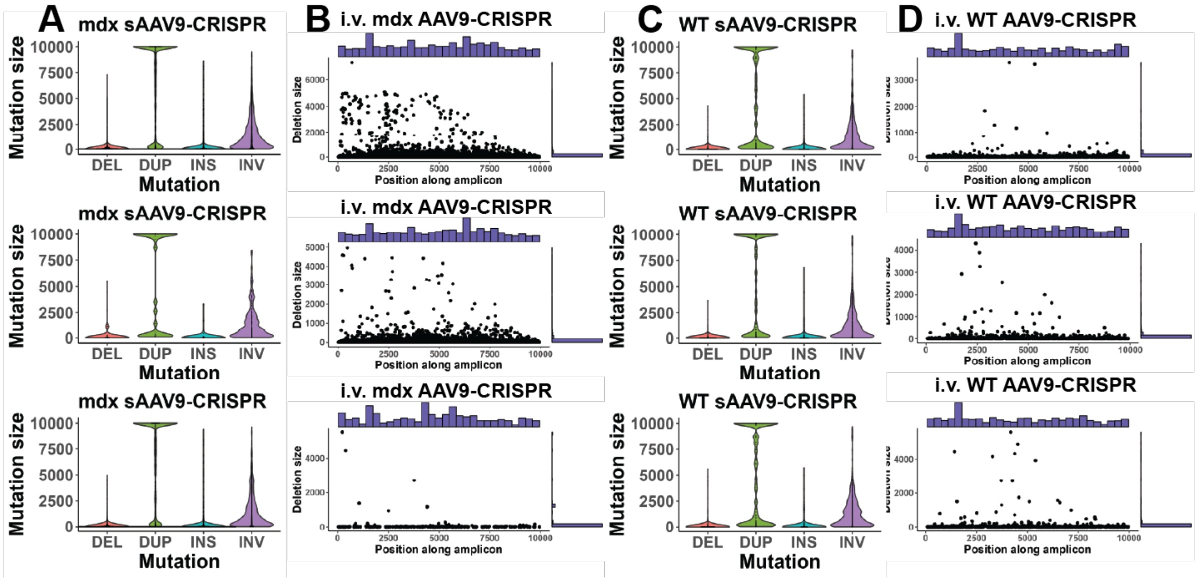


**Figure S4**. Mutation profiles following intramuscular AAV9 or AAV8 CRISPR delivery. Left panels show the distribution of deletions across the amplicon positions, with histograms showing the frequency of events. Right panels illustrate the mutation size distribution categorized by deletion (DEL), duplication (DUP), insertion (INS), and inversion (INV) for each experimental group (i.m. AAV9-CRISPR and i.m. AAV8-CRISPR in both mdx and WT models). The data highlight mutation diversity and frequency induced by CRISPR delivery systems

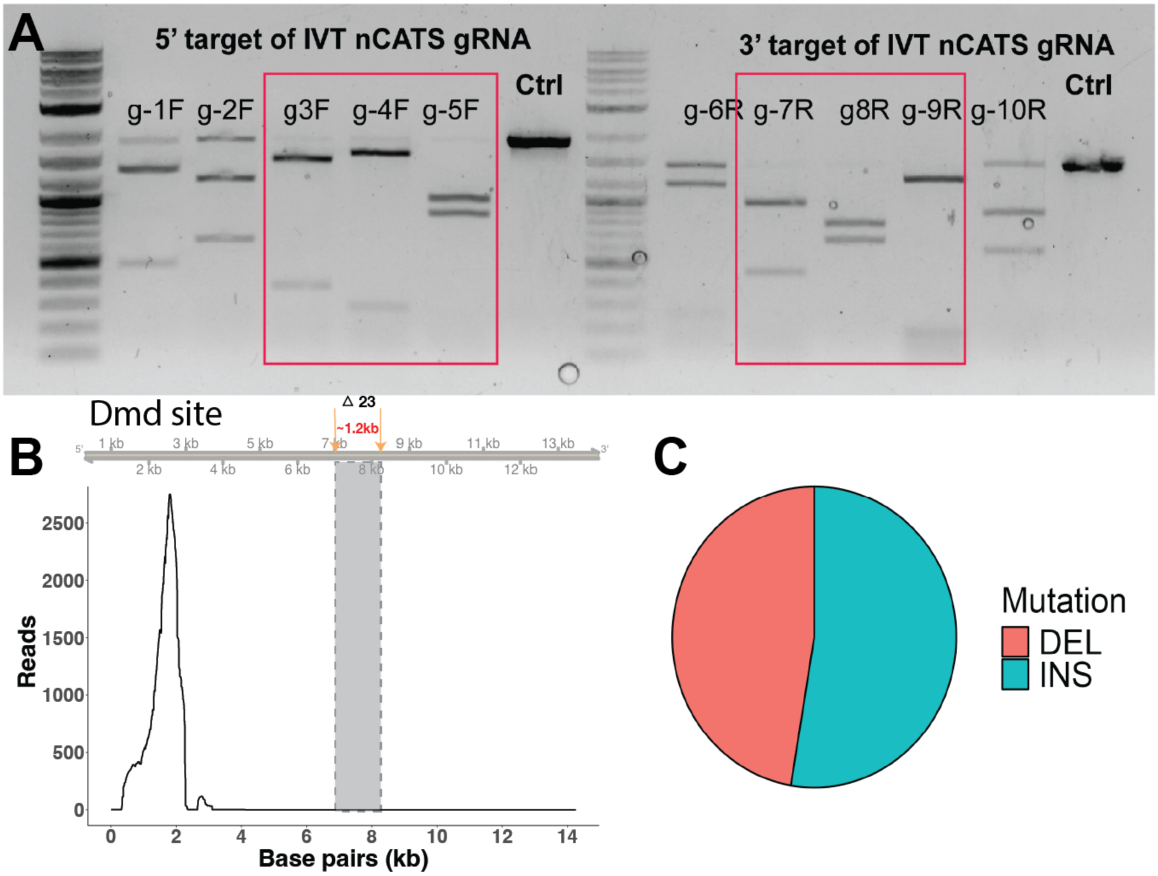


**Figure S5**. (**A**) In vitro digestion confirmation of IVT nCATS gRNA. The figure shows the cleavage efficiency of individual gRNAs targeting the 5' and 3' regions. Red squares in the gRNA lanes demonstrate near 100% efficiency cleavage of respective gRNA. (**B**) Tn5-mediated whole-genome sequencing fails to enrich at the target editing site - illustrating insufficient coverage to detect rare editing events.

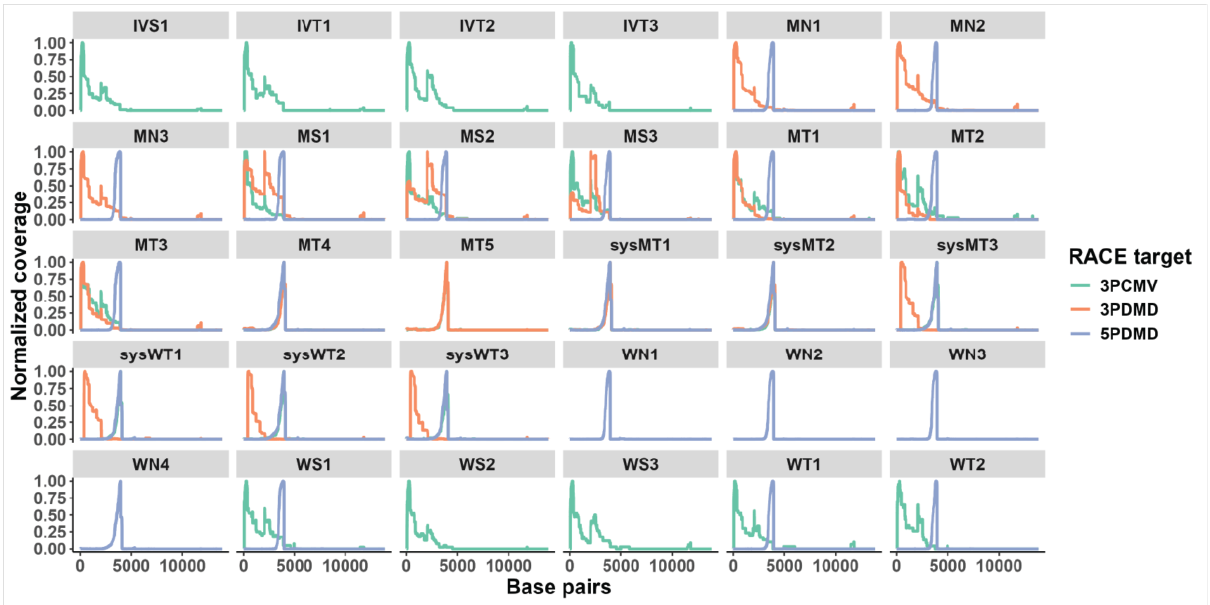


**Figure S6.** Normalized coverage of RACE targets using 3 and 5 RACE strategies across multiple experimental groups aligned to the full-length Dmd transcript. Each panel represents normalized coverage profiles for RACE targets: 3PCMV (3 RACE using CMV as gene-specific primer, green), 3PDmd (3 RACE using Dmd as gene-specific primer, orange), and 5PDmd (5 RACE using Dmd as gene-specific primer, blue). The x-axis denotes base pairs (in kb), and the y-axis represents normalized coverage. Variations in coverage across different experimental conditions highlight the efficiency and specificity of RACE primer-target interactions in detecting structural variants and transcript diversity. IV – intravenous delivery; M – mdx mice; N – non-treated; S – scrambled control; T – treated; W – wild-type.

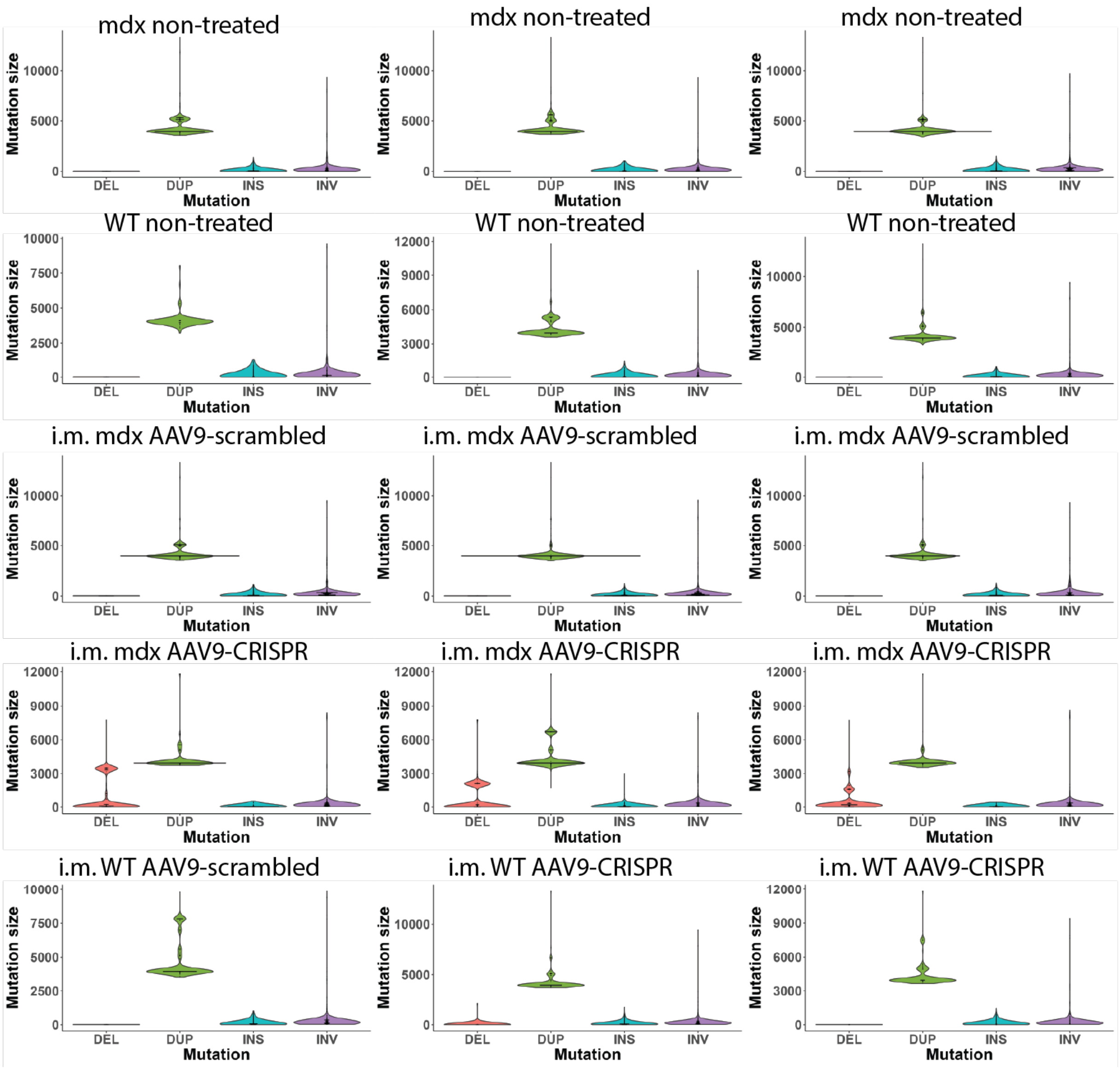


**Figure S7.** Structural variant detection using 5' RACE following intramuscular AAV9-CRISPR Treatment. The figure shows the impact of AAV9-CRISPR and scrambled control treatments on structural variants across different experimental conditions. Violin plots represent the size distribution of structural variants identified from 5′ RACE reads in mdx and wild-type (WT) mice treated with AAV9-CRISPR via intramuscular (i.m.) delivery. Detected mutation types include deletions (DEL), duplications (DUP), insertions (INS), and inversions (INV). Each panel corresponds to a different biological replicate or condition. DUPs and INSs are the most prominent mutation types across both genotypes, with generally larger mutation sizes observed in duplications.

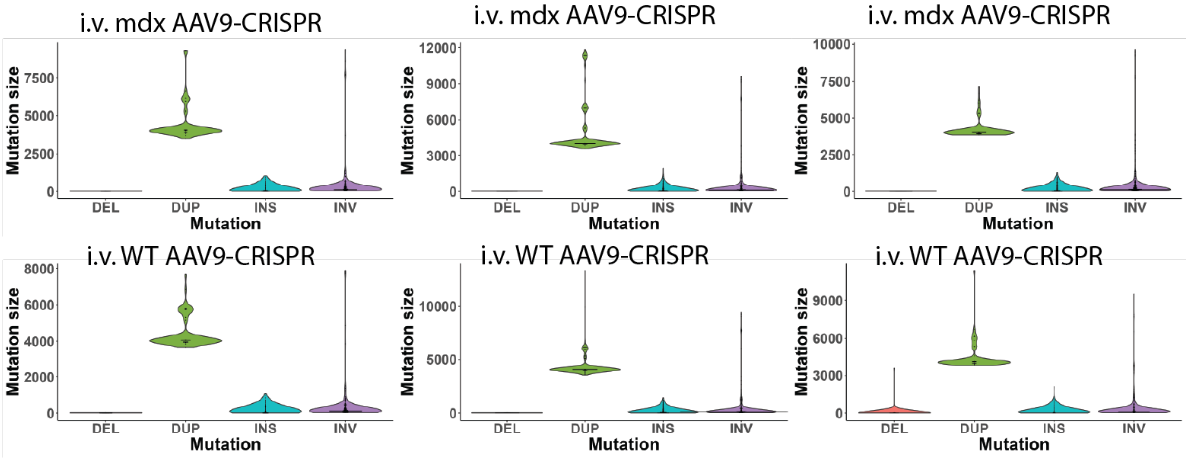


**Figure S8.** Structural variant detection using 5’ RACE following intravenous AAV9-CRISPR Treatment. The results are indistinguishable from unedited samples due to low editing efficiency in systemic treatment groups.

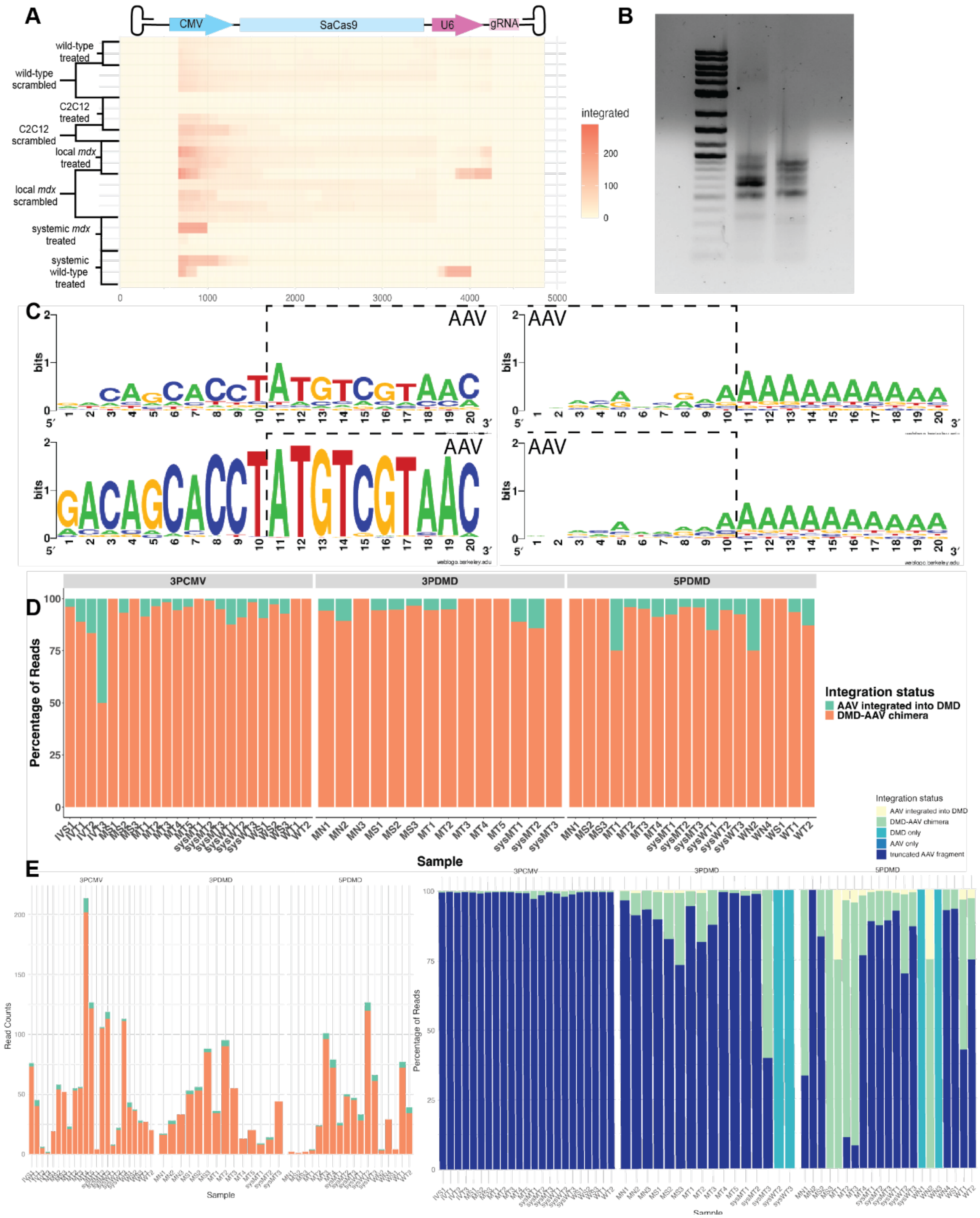


**Figure S9**: nRACE-seq reveals a diverse set of mutation events causing altered transcripts. (**A**) Heatmap of reads mapped to the full-length AAV genome that also map to the *Dmd* transcript that are indicative of fusion events. (**B**) Gel of 5' RACE products demonstrate a range of amplification products used in sequencing. (**C**) Replicates of *mdx* edited mice nRACE-seq results illustrate conserved motifs present where AAV integration either begins or terminates. The initiation site appears to be significantly more conserved than the termination site upstream of the poly A tail. (**D**) Sequence counts normalized to total sequenced reads for each enrichment technique sorted by the location of the detected AAV fusion event. Location and nature of AAV fusion events detected are dependent on the nRACE-seq technique used to enrich for transcripts. Enrichment of an unknown 3' end with primers specific for the putative CMV transcriptional start site detect the broadest range of fusion events with primarily AAV transcripts that are flanked at the 5' and 3' end by *Dmd*. (**E**) The same illustration as in D without normalization using the raw read counts. 3'RACE CMV enriching for the AAV transcript was the most successful in achieving a large number of reads. The adjacent figure illustrate the normalized full set of reads including sequencing reads that only contained AAV and *Dmd* transcripts. The majority of reads map to truncated AAV genomes, which may be indicative of alternative AAV integration sites. Further, while 3'CMV race detected the most fusion reads, the proportion of reads that exist as fusion events compared to 3' and 5' RACE amplifying for *Dmd* are lower.

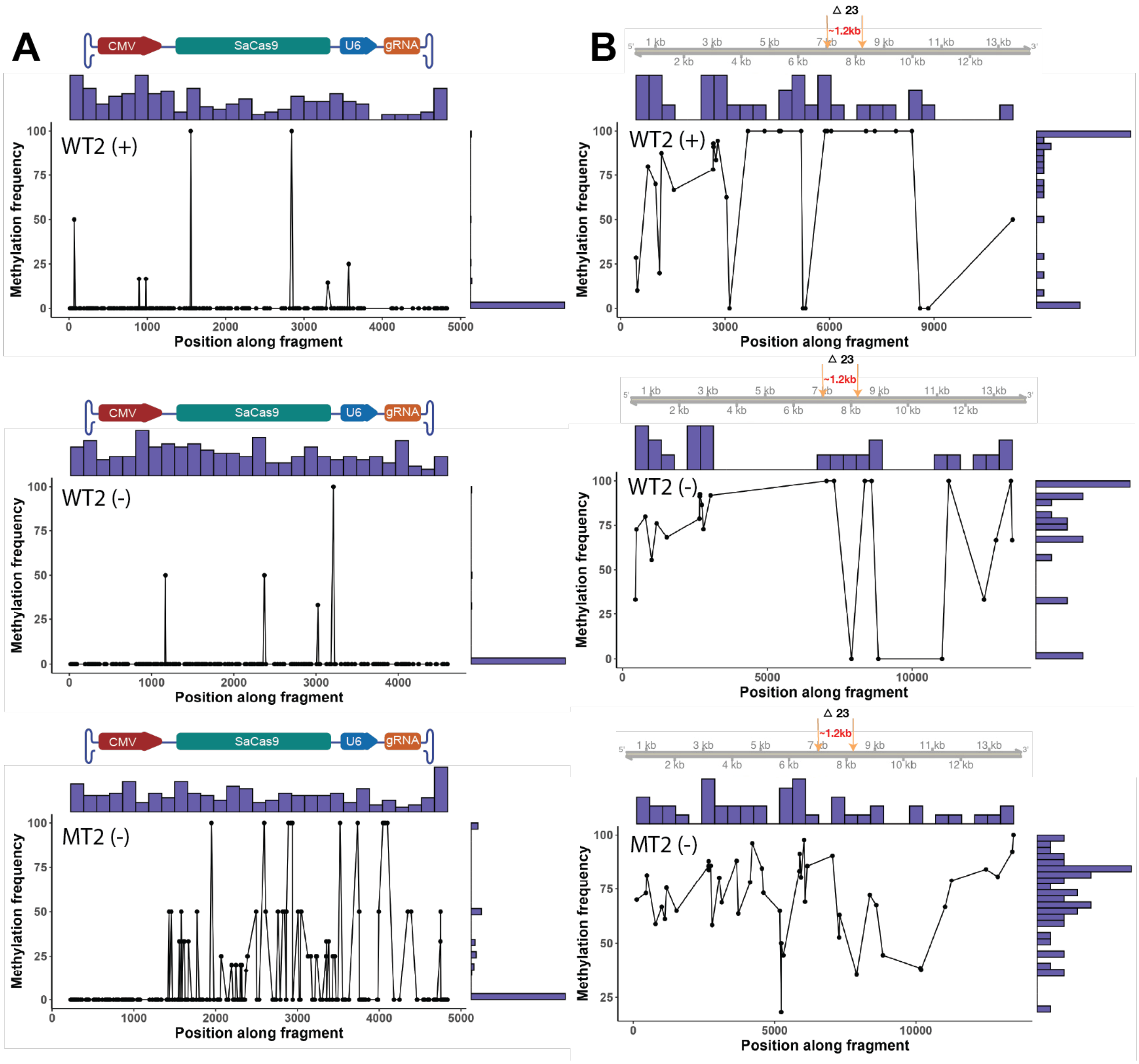


**Figure S10**: Methylation detection along AAV and *Dmd* sequences with PCR-free enrichment. (**A**) Methylation frequencies of detected integrated AAV sequences in wild-type AAV9-CRISPR-edited IM mice with whole-genome sequencing and *mdx* AAV9-CRISPR-edited IM mice with nCATS-AS, respectively. Methylation frequencies based on the forward and reverse strand are plotted separately to illustrate strand bias for detection. Due to significantly higher coverage of the target site in nCATS-AS, the resulting methylation detection is more robust from base to base. (**B**) Methylation frequencies detected along the Dmd 13kb enriched in nCATS-AS for the same sample groups as in A.

**Table S1:**

| Category | Primer Name | Sequence |
|---|---|---|
| gRNA target | Guide 1 intron 22 | TACACTAACACGCATATTTG |
| gRNA target | Guide 2 intron 23 | CATTGCATCCATGTCTGACT |
| PCR-enriched Iseq sequencing | Fwd primer intron 22 Dmd | TCGTCGGCAGCGTCAGATGTGTATAAGAGACAG-tttctgtctaaatataatatgccctgt |
| PCR-enriched Iseq sequencing | Rev primer intron 22 Dmd | GTCTCGTGGGCTCGGAGATGTGTATAAGAGACAG-gcagagcctcaaaattaaatagaag |
| PCR-enriched Iseq sequencing | Fwd primer intron 23 Dmd | TCGTCGGCAGCGTCAGATGTGTATAAGAGACAG-TGTTTCAAGAGCTCATCCTCTTTC |
| PCR-enriched Iseq sequencing | Rev primer intron 23 Dmd | GTCTCGTGGGCTCGGAGATGTGTATAAGAGACAG-AGAAAATGCAAAAGGACCCCC |
| Tn5 Sequencing | GSP for Dmd intron 22 | GTCTCGTGGGCTCGGAGATGTGTATAAGAGACAG-CTGTCTAAATATAATATGCCCTG |
| Tn5 Sequencing | GSP for Dmd intron 23 | GTCTCGTGGGCTCGGAGATGTGTATAAGAGACAGCAG-GCAGGAGCAACTTTGGA |
| Tn5 Sequencing | Universal i5 | AATGATACGGCGACCACCGAG-ATCTACACTCGTCGGCAGCGTC |
| DNA PCR amplification | Fwd primer DNA - PCR amplicon | TTATTTCTTCCAAACATATTCTCATCCACA |
| DNA PCR amplification | Rev primer DNA - PCR amplicon | TGCAAATCACCACATGTTCCAT |
| cDNA PCR amplification | Fwd primer cDNA-PCR amplicon | GATCAAAATGAAATGATGTCAAGTCTTC |
| cDNA PCR amplification | Rev primer cDNA-PCR amplicon | GATTTGTTCTATGTTCTGATCAAAGGTTTC |
| 5’ RACE and 3’ RACE | TSO | GCTAATCATTGCAAGCAGTGGTATCAACGCAGAG-TACATrGrGrG |
| 5’ RACE and 3’ RACE | TSO2 | CATTGCAAGCAGTGGTATCAAC |
| 5’ RACE and 3’ RACE | 5’ RACE GSP for Dmd | GATTTGTTCTATGTTCTGATCAAAGGTTTC |
| 5’ RACE and 3’ RACE | 3’ RACE GSP for CMV | ATGTCGTAACAACTCCGCCC |

| | | |
|---|---|---|
| 5’ RACE and 3’ RACE | TSO-specific primer | GCTAATCATTGCAAGCAGTGGT |
| 5’ RACE and 3’ RACE | Poly-T hairpin | CGTATCCAGTGCAGGGTCCGAGGTATTCGCACT-GGATACGTTTTTTTTTTTTTTTTTTT |
| 5’ RACE and 3’ RACE | Poly-T-hairpin specific primer | CAGTGCAGGGTCCGAGGTAT |
| In vitro guide RNA for nCATS | nCATS1F | TAATACGACTCACTATAGGggACTTGAGTCATGAGTGCACT-gttttagagctagaaatag |
| In vitro guide RNA for nCATS | nCATS2F | TAATACGACTCACTATAGGggTTATGCTGCACATATCCCAT-gttttagagctagaaatag |
| In vitro guide RNA for nCATS | nCATS3F | TAATACGACTCACTATAGGggATAACTTACAGCTCTTGGCA-gttttagagctagaaatag |
| In vitro guide RNA for nCATS | nCATS4F | TAATACGACTCACTATAGGggCAGGTCACAGAAATGGCACA-gttttagagctagaaatag |
| In vitro guide RNA for nCATS | nCATS5F | TAATACGACTCACTATAGGggATTCCCATAATAAATTCCAT-gttttagagctagaaatag |
| In vitro guide RNA for nCATS | nCATS6R | TAATACGACTCACTATAGGggTGTGTTGGAGGTGTGGCGTG-gttttagagctagaaatag |
| In vitro guide RNA for nCATS | nCATS7R | TAATACGACTCACTATAGGggCTAACTAAAACACTATTCCA-gttttagagctagaaatag |
| In vitro guide RNA for nCATS | nCATS8R | TAATACGACTCACTATAGGggTTAATAGAATCAATTATCCT-gttttagagctagaaatag |
| In vitro guide RNA for nCATS | nCATS9R | TAATACGACTCACTATAGGggAAAACTAATAAATCACATCC-gttttagagctagaaatag |
| In vitro guide RNA for nCATS | nCATS10R | TAATACGACTCACTATAGGggATGTGTTTAGTACTGCTG-gttttagagctagaaatag |
| In vitro guide RNA for nCATS | tracrRNA | Alt-R™ CRISPR-Cas9 tracrRNA, 20 nmol, IDT Cat. no. 1072533 |
| Surveyor in vitro digestion primers for DMD | Fwd primer 5’ region | TCAGAAGTTCAGGCCACAACT |
| Surveyor in vitro digestion primers for DMD | Rev primer 5’ region | TTGCACTGCTTTCATTGGGT |
| Surveyor in vitro digestion primers for DMD | Fwd primer 3’ region | TGCCAGTGAGTCAAAACAAATATGT |
| Surveyor in vitro digestion primers for DMD | Rev primer 3’ region | GAAGCTGACCCCTGTGGAAA |
